# Possible Cross Reactivity of Feline and White-tailed Deer Antibodies Against the SARS-CoV-2 Receptor Binding Domain

**DOI:** 10.1101/2021.12.17.473265

**Authors:** Trevor J. Hancock, Peyton Hickman, Niloo Kazerooni, Melissa Kennedy, Stephen A. Kania, Michelle Dennis, Nicole Szafranski, Richard Gerhold, Chunlei Su, Tom Masi, Stephen Smith, Tim E. Sparer

## Abstract

In late 2019, a novel coronavirus began circulating within humans in central China. It was designated SARS-CoV-2 because of its genetic similarities to the 2003 SARS coronavirus (SARS-CoV). Now that SARS-CoV-2 has spread worldwide, there is a risk of it establishing new animal reservoirs and recombination with native circulating coronaviruses. To screen local animal populations in the United States for exposure to SARS-like coronaviruses, we developed a serological assay using the receptor binding domain (RBD) from SARS-CoV-2. SARS-CoV-2’s RBD is antigenically distinct from common human and animal coronaviruses allowing us to identify animals previously infected with SARS-CoV or SARS-CoV-2. Using an indirect ELISA for SARS-CoV-2’s RBD, we screened serum from wild and domestic animals for the presence of antibodies against SARS-CoV-2’s RBD. Surprisingly pre-pandemic feline serum samples submitted to the University of Tennessee Veterinary Hospital were ∼50% positive for anti-SARS RBD antibodies. Some of these samples were serologically negative for feline coronavirus (FCoV), raising the question of the etiological agent generating anti-SARS-CoV-2 RBD cross-reactivity. We also identified several white-tailed deer from South Carolina with anti-SARS-CoV-2 antibodies. These results are intriguing as cross-reactive antibodies towards SARS-CoV-2 RBD have not been reported to date. The etiological agent responsible for seropositivity was not readily apparent, but finding seropositive cats prior to the current SARS-CoV-2 pandemic highlights our lack of information about circulating coronaviruses in other species.

**Importance:** We report cross-reactive antibodies from pre-pandemic cats and post-pandemic South Carolina white-tailed deer that are specific for that SARS-CoV RBD. There are several potential explanations for this cross-reactivity, each with important implications to coronavirus disease surveillance. Perhaps the most intriguing possibility is the existence and transmission of an etiological agent (such as another coronavirus) with similarity to SARS-CoV-2’s RBD region. However, we lack conclusive evidence of pre-pandemic transmission of a SARS-like virus. Our findings provide impetus for the adoption of a One Health Initiative focusing on infectious disease surveillance of multiple animal species to predict the next zoonotic transmission to humans and future pandemics.

## Introduction

Severe acute respiratory syndrome coronavirus 2 (SARS-CoV-2) is an emergent zoonotic beta-coronavirus initially identified in late 2019 after human-to-human transmission within central China [1]. By early 2020, the virus had caused a pandemic infecting millions of people and continues to circulate throughout the world. Like other human coronaviruses it is spread *via* aerosolized particles, leading to respiratory infections [1, 2]. Infected individuals develop a range of symptoms from mild/asymptomatic infection to severe pneumonia-like disease (i.e., coronavirus disease (COVID)) [2]. Sequence analysis of known SARS coronaviruses points to a bat origin with probable intermediate hosts prior to human adaptation [3-5]. However, the exact intermediate host and factors that led to its zoonosis and establishment within humans are under investigation.

Secretion of SARS-CoV-2 is thought to be primarily *via* aerosolized particles with high viral loads in the lungs and nasopharyngeal secretions of infected individuals [6, 7]. However, both viral RNA and infectious particles have been detected in fecal samples of acutely infected individuals. In the original SARS-CoV outbreak, there was documented fecal-oral transmission of infection [6, 8-12]. Fecal to oral spread and shedding is a common route of transmission of other animal coronaviruses. Oropharyngeal viral RNA shedding of SARS-CoV-2 in humans lasts for ∼17 days on average but persists up to 60-120 days in the respiratory tract and stool [13]. Similarly, oropharyngeal secretion of infectious SARS-CoV-2 in cats appears to cease by 5-10 days post infection (dpi) [14]. Infected felids shed SARS-CoV-2 viral RNA in their feces for at least 5 dpi, but whether that represents infectious virus, or a potential route of transmission is yet to be demonstrated [15].

Due to the multiple routes of spread and close contact with other species, transmission of SARS-CoV-2 from humans to animals is plausible [16]. Human-to-animal and animal-to-animal transmission of SARS-CoV-2 has been documented or experimentally demonstrated. Companion animals such as cats and dogs are susceptible to experimental as well as natural infection from COVID-positive owners [14, 17-21]. In addition, susceptible animals are capable of transmitting infection to cohoused animals [14, 22]. In the case of minks, transmission from humans-to-minks and back to humans was demonstrated [23]. This is similar to a situation at the Amoy Garden complex during the original SARS-CoV outbreak where animal-to-human transmission occurred when an animal vector potentially contracted and spread SARS-CoV throughout the complex [24-26]. Human transmission of SARS-CoV-2 to companion animals opens up the potential for spillover into wild animal populations. Indeed, transmission from humans to deer within North America has been proposed, as post-pandemic deer have been demonstrated seropositive in multiple states and SARS-CoV-2 genome sequenced from lymph nodes [27-29]. Human to animal transmission could contribute to the spread of SARS-like coronaviruses and the establishment of new reservoirs for recombination and the generation of future novel coronavirus outbreaks.

Infected humans and animals mount humoral responses to SARS-CoV-2 [13, 14, 30-32]. In humans, SARS-CoV-2 antibodies arise within 5-14 days post-infection/symptom onset and peak around 17-20 dpi [13, 31, 33]. For cats experimentally inoculated or naturally exposed to SARS-CoV-2, detectable antibody titers appeared by 7-14 days post-infection peaking ∼21 dpi [14]. This matches anti-FCoV responses where high antibody levels can arise within ∼9 dpi [34-36]. Immunity to coronaviruses in cats is typically short-lived, with the average FCoV humoral responses lasting several months to 2 years [37]. Anti-SARS-CoV-2 RBD responses in seropositive cats had similar declines in antibody titers only lasting around 4-5 months [38]. However, humans infected with the initial SARS-CoV mounted robust responses detectable 1-2 years post exposure [39-41]. The duration of anti-SARS-CoV-2 antibody responses is the subject ongoing research, but natural exposure is unlikely to induce long-term or lifelong immunity/seropositivity [42].

**Table 1:**
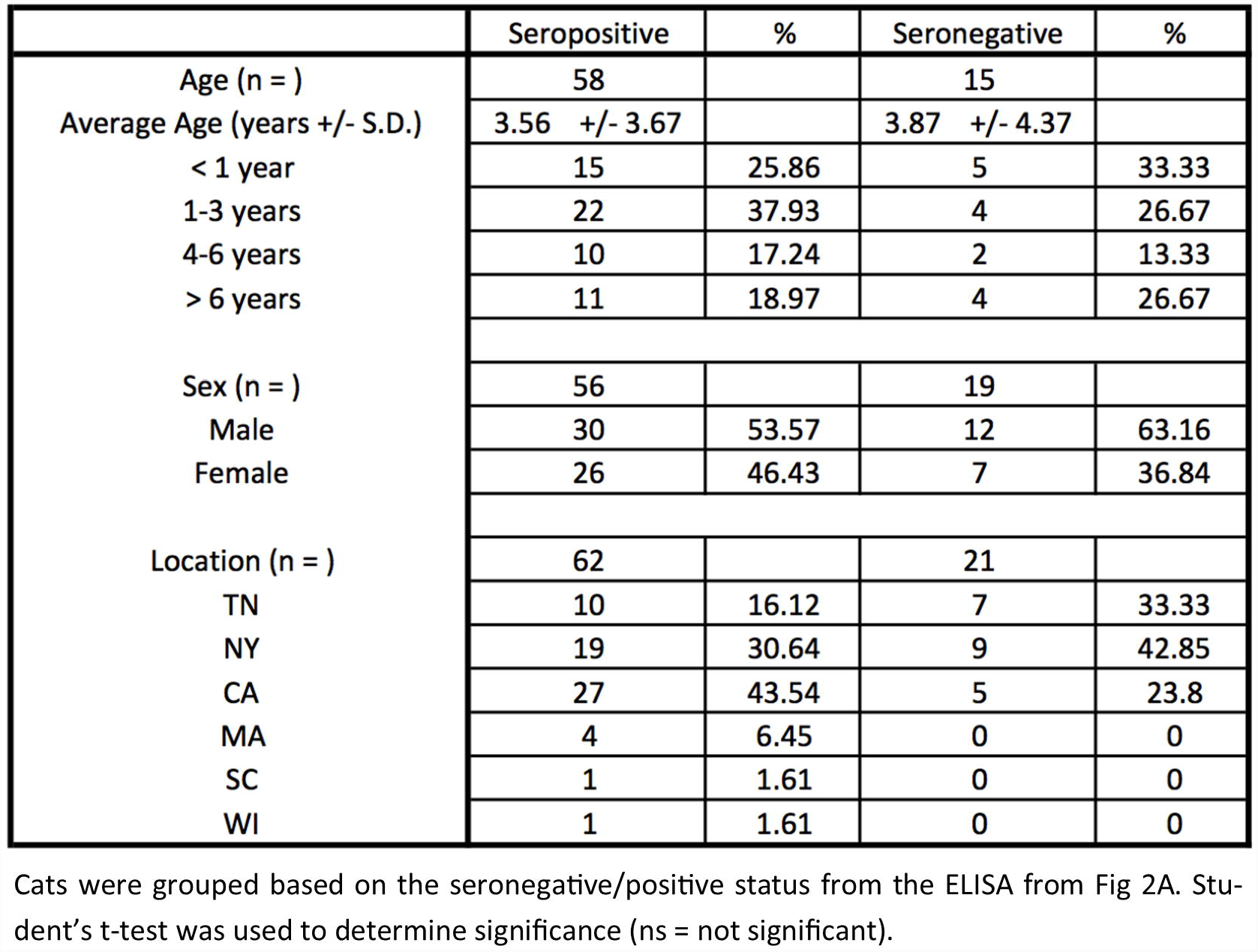
Characteristics of Feline Samples.

**Table 2:** Fecal Coronavirus Screening Information.

| Sample | RdRp1 | RdRp2 | Spike1 | Spike2 | Spike3 | Helicase | Location | Accession #'s |
| --- | --- | --- | --- | --- | --- | --- | --- | --- |
| CP-20-01 |  |  |  |  |  |  | Jefferson County, TN, USA | MZ220722-MZ220724 |
| CP-20-02 |  |  |  |  |  |  | Washington County, TN, USA | MZ220725-MZ220727 |
| CP-20-03 |  |  |  |  |  |  | TN, USA | N/A |
| CP-20-04 |  |  |  |  |  |  | Monroe County, TN, USA | MZ220728-MZ220730 |
| CP-20-05 |  |  |  |  |  |  | TN, USA | N/A |
| CP-20-06 |  |  |  |  |  |  | Sullivan County, TN, USA | N/A |
| CP-20-07 |  |  |  |  |  |  | Grainger County, TN, USA | MZ220731-MZ220734 |
| CP-20-08 |  |  |  |  |  |  | Monroe County, TN, USA | MZ220735 |
| CP-20-09 |  |  |  |  |  |  | Sullivan County, TN, USA | MZ220736-MZ220738 |
| CP-20-10 |  |  |  |  |  |  | Washington County, TN, USA | N/A |
| CP-20-11 |  |  |  |  |  |  | Jefferson County, TN, USA | N/A |
| CP-20-12 |  |  |  |  |  |  | Monroe County, TN, USA | MZ220739-MZ220742 |
| CP-20-13 |  |  |  |  |  |  | Grainger County, TN, USA | MZ220743-MZ220745 |
| CP-20-14 |  |  |  |  |  |  | Jefferson County, TN, USA | N/A |
| CP-20-15 |  |  |  |  |  |  | Sullivan County, TN, USA | N/A |
| CP-20-16 |  |  |  |  |  |  | Sevier County, TN, USA | N/A |
| CP-20-17 |  |  |  |  |  |  | Cocke County, TN, USA | N/A |
| CP-20-18 |  |  |  |  |  |  | Hamblin County, TN, USA | N/A |
| CP-20-19 |  |  |  |  |  |  | Washington/Sullivan County, TN, USA | MZ220746-MZ220747 |
| CP-20-20 |  |  |  |  |  |  | Hamblin County, TN, USA | MZ220748 |
| CP-20-21 |  |  |  |  |  |  | Washington/Sullivan County, TN, USA | N/A |
| CP-20-22 |  |  |  |  |  |  | Grainger County, TN, USA | N/A |
| CP-20-23 |  |  |  |  |  |  | Jefferson County, TN, USA | MZ220749-MZ220750 |
| CP-20-24 |  |  |  |  |  |  | Jefferson County, TN, USA | N/A |
| CP-20-25 |  |  |  |  |  |  | Grainger County, TN, USA | N/A |
| CP-20-26 |  |  |  |  |  |  | Grainger County, TN, USA | MZ220751-MZ220755 |
| CP-20-27 |  |  |  |  |  |  | Grainger County, TN, USA | MZ220756-MZ220759 |
| CP-20-28 |  |  |  |  |  |  | Washington/Sullivan County, TN, USA | MZ220760 |
| CP-20-29 |  |  |  |  |  |  | TN, USA | N/A |
| CP-20-30 |  |  |  |  |  |  | Grainger County, TN, USA | MZ220761-MZ220762 |

Sample IDs were assigned to de-identified fecal samples from healthy cats. RNA was extracted from each and PCR screened for amplification of 6 coronavirus loci across 3 conserved genes (RdRp, spike, and helicase). Positive amplification and sequencing is shown by a black shaded box. 15/30 samples were positive for at least one coronavirus loci. Animals originated from multiple East Tennessee counties, and, when known, exact county of origin is listed. Accession numbers for each sample are shown to the right and consist of each sequenced loci.

Major antigenic targets for SARS-CoV-2 infected individuals are the nucleocapsid, which is one of the most abundantly produced viral proteins [43], and spike protein, which is responsible for viral entry [44]. The spike has high immunogenicity and diverges from other coronaviruses [32, 44, 45]. Spike is composed of two subunits (S1/S2). The S1 subunit contains the receptor binding domain (RBD) responsible for binding to host ACE-2 and determining tropism/entry, while the S2 domain contains the fusogenic region of the spike [44, 45]. SARS coronaviruses share very low similarity to other coronaviruses within the spike protein [32], but antibodies against the S2 subunit can cross-react with common human coronaviruses [46-48]. Cross-reactivity of the S1 subunit occurs at very low rates. Within the S1 region, the RBD is highly immunogenic and unique to SARS-CoV-2 [32, 49]. Serum from humans infected with common human coronaviruses such as OC43, NL63, and 229E failed to recognize the RBD from SARS-CoV-2 [32, 46, 49]. Animals infected or immunized with other coronaviruses similarly fail to generate cross-reactive antibodies against SARS-CoV-2’s RBD [32]. For infected cats, SARS-CoV-2 seroconversion was not impacted by pre-existing immunity against feline coronavirus (FCoV), an alpha-coronavirus with limited similarity to SARS-CoV-2 [38]. Collectively, seropositivity against the RBD of SARS-CoV-2 is a specific marker of SARS-CoV-2 exposure and has led several groups to create highly specific indirect ELISAs against SARS-CoV-2’s RBD to screen for SARS-CoV-2 exposure [30, 32, 33]. A final consideration of antibodies targeting the RBD is they could be either neutralizing or non-neutralizing [33, 50-53]. This may explain why serum from humans and animals exposed to the original SARS-CoV were able to recognize the spike and RBD of SARS-CoV-2 while their cross-neutralization potential was variable [54, 55].

Despite limited similarity in the spike protein of SARS-CoV-2 vs common circulating coronaviruses, there are reports of pre-pandemic, pre-existing SARS-CoV-2 reactive serum in humans [48, 49, 54]. These cross-reactive antibodies represent a rare response to common human coronaviruses within conserved epitopes of SARS-CoV-2’s spike protein (usually in the S2 region) with reports of ∼0.6% prevalence of pre-existing anti-RBD responses [46, 48, 49]. Although there is increasing evidence for earlier timelines of SARS-CoV-2 spread among humans, pre-existing seropositivity among other species has not been reported [38, 56-58]. Indeed, even within central China, researchers failed to find evidence of SARS-CoV-2 exposure prior to the pandemic [38, 56, 58].

As SARS-CoV-2 spreads and encounter’s new species, there is a need for monitoring local populations for SARS-CoV-2 transmission and the potential establishment of local reservoirs. Currently, we have a limited understanding of coronavirus reservoirs, spread, and recombination among diverse species. The original SARS outbreak in 2003 was a harbinger of the potential risk of crossover coronaviruses. At that time, animal coronavirus surveillance was a high priority. Unfortunately, this investment was not sustained. Our aim was to address whether SARS-CoV-2 is being introduced into companion animals of North America by tracking seroconversion using an in-house indirect ELISA against the RBD of SARS-CoV-2. We chose to focus on companion animals (i.e., cats and dogs) as they represent a significant source of human-animal interactions with potential for contact and further spillover into wild animal populations. Surprisingly, we found evidence of anti-RBD seropositive animals pre-dating the pandemic by several months to years. Our study provides evidence for the existence and prevalence of SARS-CoV-2 serum reactivity prior to the current pandemic.

## Materials and methods

### Recombinant RBD production and purification

Recombinant RBD production has been previously published [56]. Our lab deviated from the prior published method to utilize equipment readily available. Briefly, the plasmid containing the RBD of SARS-CoV-2 was produced under federal contract HHSN272201400008C and obtained through BEI Resources, NIAID, NIH. Vector pCAGGS contains the SARS-related coronavirus 2, Wuhan-Hu-1 spike glycoprotein RBD, NR-52309. To produce recombinant RBD, the pCAGGS-RBD plasmid was transfected into ∼5×10^7^ adherent HEK-293/T17 cells (ATCC CRL-11268) in a T-175 using PEI (Polyethylenimine, linear 25,000 m.w.(Polysciences, Warrington, PA, USA). Plasmid was mixed at a 1:3 ratio with PEI (20ug of plasmid : 60ug PEI for a T-175 transfection) in 1 mL serum-free DMEM for 30 minutes at room temperature. Media was aspirated and the transfection mixture was added to 14mL fresh growth media and placed onto cells. Three to four hours post-transfection, media was changed and replaced with DMEM containing either 2% or 5% Fetal Bovine Serum (Hyclone FetalClone III, Cytiva Life Sciences, USA). Maintenance in a lower serum prevents overgrowth. However, we found higher protein yields when supplemented with 5% FBS. Transfection efficiency was nearly 100% as assessed by GFP-positive transfected cells in a control flask.

Supernatants from transfected HEK-293 T17 cells were collected into 50mL conical tubes and frozen at days 3 and 6 post-transfection. Pooled supernatants were thawed and incubated with Ni-NTA (Ni-NTA Agarose, Qiagen, Germany) resin with gentle rocking overnight. The resin was spun down at >3400x g in a swing-bucket Sorvall RT centrifuge for 10 minutes at 4^0^C. Ni-NTA resin was resuspended in 1mL wash buffer (20mM imidazole, 5mM NaH_2_PO_4_.H_2_O, 0.3M NaCl in H_2_0), transferred to a 2mL microcentrifuge tube, gently rocked for 10 minutes at room temperature, spun, and resuspended in fresh buffer. Resin was washed between 3-7 times until OD_230_ was ≤ wash buffer. Once the supernatant OD dropped sufficiently, 1mL elution buffer (235mM Imidazole, 5mM NaH_2_PO_4_.H_2_O, 0.3M NaCl in H_2_0) was added to elute the RBD from the nickel resin. Eluate was rocked for 10 minutes at room temp and then centrifuged. Two elution steps were performed with a third final elution using 0.5M imidazole. Protein concentration was determined by standard curve analysis of a silver-stained (Pierce Silver Stain Kit, Thermo Scientific, USA) 12% SDS-PAGE gel using a standard curve of BSA (bovine serum albumin). Analysis was performed using Image Studio Lite ver. 5.3 (Li-Cor Biosciences, Lincoln, NE, USA).

### Serum and plasma samples

Previously collected pre-SARS-CoV-2 de-identified human serum samples were kindly donated by Dr. Jon Wall and Steve Foster (University of Tennessee Medical Center, Knoxville, Tennessee, USA). De-identified COVID-positive plasma samples were donated from: MEDIC Regional Blood Center (Knoxville, Tennessee, USA) and Dr. Mark Slifka (Oregon Health Sciences University, Portland, Oregon, USA). The following reagents were obtained through BEI Resources, NIAID, NIH: Human Plasma, Sample ID WU353-073, NR-53643; WU353-074, NR-53644; WU353-075, NR-53645; WU353-076, NR-53646; WU353-076, NR-53647, were contributed by Ali Ellebedy, Ph.D., Washington University School of Medicine, St. Louis, Missouri, USA. The following reagents were obtained through BEI Resources, NIAID, NIH: polyclonal anti-feline infectious peritonitis virus, 79-1146 (antiserum, guinea pig), NR-2518; polyclonal anti-canine coronavirus, UCD1 (antiserum, guinea pig), NR-2727; polyclonal anti-bovine coronavirus, Mebus (antiserum, guinea pig), NR-455; polyclonal anti-porcine respiratory coronavirus, ISU-1 (antiserum, guinea pig), NR-459; polyclonal anti-turkey coronavirus, Indiana (antiserum, guinea pig), NR-9465. Client-owned canine and feline serum samples were submitted to the University of Tennessee Veterinary Hospital for routine animal testing. Canine samples (n=36) were collected post-pandemic from local, East Tennessee animals. All feline samples were pre-pandemic. Samples from East Tennessee feral cats (n=36) were collected from 2007-2012. Client-owned feline samples (n=92) were collected nationwide as part of clinical diagnostic testing (see Table 1). Twenty cat samples were grouped into FCoV positive and negative groups based on feline infectious peritonitis (FIP) serology using an immunofluorescence assay (IFA) against FIP serotypes I and II, as well as TGEV (transmissible gastroenteritis virus) (VMRD, Pullman, WA, USA). Normal cat serum was purchased from Jackson ImmunoResearch (West Grove, PA, USA). Tennessee-resident cows (n=33) and tigers (n=9) were collected pre-pandemic for routine diagnostic testing. Post-mortem, post-pandemic samples were collected from East Tennessee elk (n=12) and South Carolina white-tailed deer (n=22).

### Anti-RBD ELISA

Anti-RBD ELISA was based on the published protocol by Amanat et al. and Stadlbauer et al. [30, 59]. Purified RBD was diluted to 2ug/mL in PBS and 50uL was placed into each well of a 96 well plate (Immulon 4HBX, Thermo Fisher, USA) and allowed to incubate overnight at 4^0^C. Unbound RBD was removed and wells were washed 3x with PBS-T (PBS with 0.1% Tween-20). Rinsed wells were blocked with 5% milk in PBS for 2 hours at room temp. Block was removed and serum or plasma samples were added at 1:50 dilution for the initial screen in PBS with 1% milk and incubated at room temp for 1 hour. After 1 hour, wells were washed 3x with PBS-T and a secondary antibody for that species was added (i.e., HRP goat-anti-human IgG, Rockland Immunochemicals, Pottstown, PA, USA; HRP goat-anti-dog IgG, Bethyl Laboratories, Montgomery, TX, USA; HRP goat-anti-cat IgG, Invitrogen, Waltham, MA, USA; HRP goat-anti-guinea pig IgG, Life Technologies Corp, Carlsbad, CA, USA; HRP rabbit-anti-deer IgG, KPL, Gaithersburg, MD, USA; HRP sheep-anti-cow IgG, Bethyl Laboratories, Montgomery, TX, USA) at dilutions of 1:10,000 (anti-human, cat, dog, tiger) or 1:250 (anti-cow, deer, elk) in PBS with 1% milk. Optimal secondary antibody concentrations were determined by titration on either 5% milk (negative control) or 1:50 dilution of that species serum (positive control). Secondary antibodies were allowed to incubate for 1 hour at room temperature before being washed 3x with PBS-T. ELISA was developed with 50uL TMB (1-Step Ultra TMB-ELISA, Thermo Fisher, Waltham, MA, USA) for 10 minutes. Reactions were stopped by the addition of 2M sulfuric acid and read using a BioTek Synergy 2 or Synergy H1 plate reader set at 450nm (BioTek, Winooski, VT, USA). Receiver operator curve (ROC) analysis was performed to the determine the appropriate threshold to yield 100% specificity of ELISAs performed at a 1:50 dilution. For titrations of seropositive and seronegative samples, threshold values for each dilution were calculated as the average of negative samples plus 3 times the standard deviation. Titrated samples were initially diluted 1:100 and then serially diluted 1:3 (final dilution of 1:8100). Serum dilutions were made in PBS with 1% milk and added to RBD-coated and blocked wells. OD_450_ values for each species and titrations were graphed in GraphPad Prism ver. 8 (GraphPad Software, San Diego, CA, USA).

### Western and dot blots

For dot blots, 5-10uL of sample was applied directly onto nitrocellulose membranes and allowed to dry. Western blots were loaded with 30uL (∼3ug) of purified recombinant RBD, resolved in a 12% SDS-PAGE gel and transferred to a nitrocellulose membrane. Blots were blocked overnight at 4^0^C with 5% milk in PBS. Mouse anti-6His-HRP conjugated antibody (1:5,000) (Proteintech, Rosemont, IL, USA) or polyclonal serum samples (1:20) were incubated with the blots at room temperature for 2 hours and subsequently washed 2x with TBS-T (tris buffered saline with 0.1% Tween-20). For polyclonal serum, species specific HRP anti-IgG antibodies (1:5,000 dilution) were incubated for an additional 2 hours and washed 2x as above. Chemiluminescent substrate (Pierce SuperSignal West Pico PLUS, Thermo Fisher, USA) was added and luminescence was detected using BioRad ChemiDoc (Bio-Rad, Hercules, CA, USA).

### Neutralization assays

Serum samples were screened for neutralization using LEGENDplex SARS-CoV-2 neutralizing antibody assay (BioLegend, San Diego, CA, USA) following manufacturer recommendations. Briefly, serum was diluted 1:100 and incubated with biotinylated SARS-CoV-2 S1 subunit containing the RBD and human ACE-2 (hACE-2) conjugated to fluorescent beads. Streptavidin-PE (phycoerythrin) was added to detect SARS-CoV-2 S1 subunit bound to beads/hACE-2. Binding/PE levels were detected *via* a BD LSR-II equipped with 488 and 633nm lasers (Becton Dickinson, Franklin Lakes, NJ, USA). Data was analyzed using BioLegend LEGENDplex Data Analysis Software. Mean fluorescent intensity (MFI) was normalized and graphed in GraphPad prism v9 (GraphPad Software, San Diego, CA, USA).

### Fecal coronavirus PCR screen and sequence alignments

De-identified fecal samples from thirty healthy East Tennessee cats were collected and stored at -80°C. Samples were resuspended in PBS to yield a 10% solution and centrifuged to clarify. Fecal RNA was extracted using a Qiagen viral RNA extraction kit (Qiagen, Hilden, Germany) and RNA was reverse transcribed using Verso cDNA kit with random hexamers and RT enhancer (Thermo Fisher, Waltham, MA, USA). PCR amplification of conserved coronavirus regions using previously reported primer pairs was used to screen the cDNA [60]. PCR amplicons were visualized on a 1% agarose gel and positive PCR samples were Sanger dideoxy sequenced. Sequences were viewed using 4Peaks software (Nucleobytes, Amsterdam, Netherlands). Sequences from fecal samples were Clustal W aligned to common coronaviruses, trimmed and phylogenetic trees were generated using Maximum-Likelihood method for each positive loci using MEGA X [61, 62]. For phylogenetic trees using multiple loci, aligned and trimmed sequences for each loci were concatenated together prior to Maximum-Likelihood tree construction. Phylogenetic trees were tested by bootstrap testing with 1000 iterations. Common coronavirus sequences for ORF1ab (RdRp and helicase loci) and spike were obtained from the following: porcine coronavirus HKU15 (NC039208), SARS-CoV-2 (MN988668), SARS-CoV (NC004718), porcine respiratory coronavirus/PRCoV (KY406735), human coronavirus OC43 (NC006213), MERS-CoV (NC038294), feline coronavirus/FCoV (NC002306), canine respiratory coronavirus/CRCoV (KX432213), canine coronavirus/CCoV (JQ404410), bovine coronavirus/BCoV (NC003045), avian coronavirus (NC048214), human coronavirus 229E (NC002645), transmissible gastroenteritis virus/TGEV (NC038861), murine hepatitis virus/MHV (NC048217), human coronavirus NL63 (NC005831), feline coronavirus strain UU8 (FJ938055), feline coronavirus strain UU19 (HQ392470), feline coronavirus strain Black (EU186072), feline coronavirus strain RM (FJ938051), feline coronavirus strain Felix (MG893511). All generated sequences were deposited in GenBank under accession numbers: MZ220722 through MZ220762 (sample ID, positive loci, animal location, and accession numbers are shown in Table 2).

### Statistics

All graphs and statistical analysis were performed in GraphPad Prism ver. 9 (GraphPad Software, San Diego, CA, USA). ELISA OD_450_ results were normalized to an inter-plate replicate run with all assays. Student’s one-tailed t-tests with Welch’s correction and one-way ANOVA with multiple comparisons tests were performed on ELISA results and documented in the respective figure legends. Descriptive statistics were provided for each ELISA group (mean, median, and quartiles). Receiver operator curve (ROC) analysis was performed to determine appropriate threshold values for human, cat, and deer serum samples. Area under the curve was calculated for each titrated ELISA sample and graphed. Neutralization data was normalized with negative control group (normal cat serum) representing 100% MFI.

Data will be made publicly available upon publication and upon request for peer review.

## Results

We developed an in-house ELISA to serologically screen companion animals based on a protocol developed at Mt. Sinai [30, 59]. To examine cross-reactivity of our in-house anti-SARS-CoV-2 RBD indirect ELISA, we used polyclonal guinea pig serum raised against different animal coronaviruses (Fig 1A). Consistent with previous reports, no cross-reactive antibodies for any of the common coronaviruses were found [18, 32, 38, 56]. Only antibodies from SARS-CoV or SARS-CoV-2 infected individuals reacted (Fig 1A, 1B). Human serum collected from individuals prior to the SARS-CoV-2 pandemic or plasma from recovered SARS-CoV-2 donors were used to validate our ELISA screen (Fig 1B). ROC analysis determined the positive cutoff threshold, using a value that gave highest specificity and sensitivity with pre-pandemic human serum and serum from confirmed SARS-CoV-2 infected individuals. ROC analysis was in agreement with the commonly used threshold determination method of three standard deviations above the mean negative value. Our assay based on RBD screening showed high sensitivity (96.96%) and specificity (95.45%) with 66 SARS-CoV-2 samples and 22 pre-SARS-CoV-2 samples (Fig 1B). While Stadlbauer et al. performed two diagnostic ELISAs, one with RBD and the other with full-length spike, our results using only the RBD-based screen are in good agreement with their published data. Others have also demonstrated the accuracy of an RBD-only based ELISA [33, 38]. A western blot using an anti-6His antibody (Fig 1C) shows the expected size of purified RBD with a single band ∼32kDa. This shows that our isolated RBD is the correct size and runs as a monomer. Silver stain of the same affinity-purified SARS-CoV-2 RBD demonstrates relative purity (Fig 1D). However, there are co-purified proteins present at lower levels.

**Figure 1:**
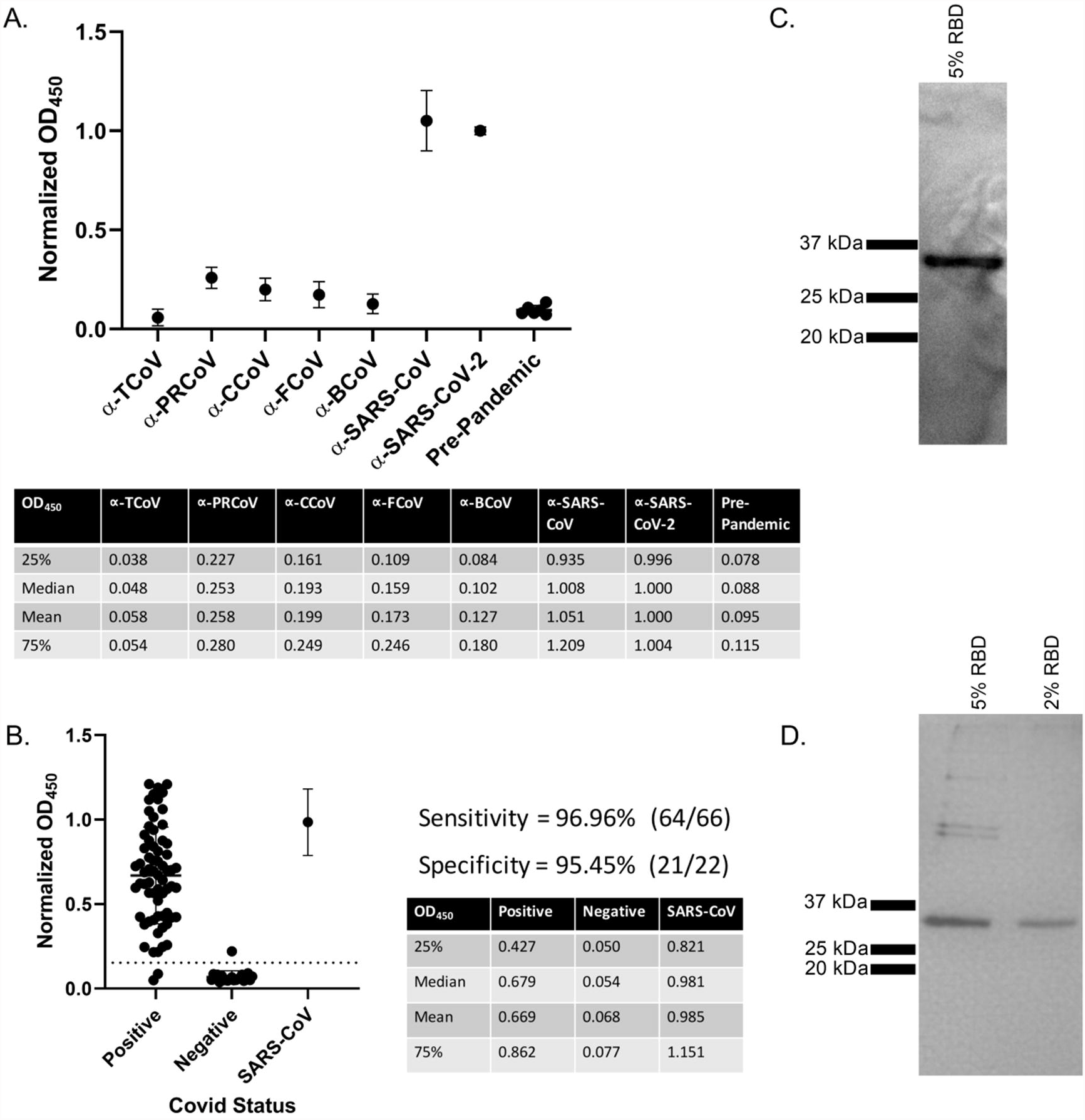
Anti-SARS-CoV-2 ELISA Sensitivity and Specificity. (A) Cross reactivity of anti-CoV antibodies against SARS-CoV-2 RBD. Polyclonal sera from guinea pigs immunized with common animal coronaviruses (turkey coronavirus, TCoV; porcine respiratory coronavirus, PRCoV; canine coronavirus, CCoV; feline coronavirus, FCoV; bovine coronavirus, BCoV) was used in a SARS-CoV-2 RBD indirect ELISA. Positive samples consisted of polyclonal serum from a SARS-CoV-2 infected patient and a monoclonal antibody to SARS-CoV (CR3022). The negative control group was comprised of pre-pandemic human serum. Secondary antibodies were either anti-human IgG (1:10,000) (Rockland Immunochemicals, USA) or anti-guinea pig IgG (1:10,000) (Life Technologies Corp, USA). Bars represent mean and standard deviation (n>3 for all samples). (B) ELISA validation using 66 human Covid-positive plasma and 22 negative serum samples. Human antibodies against the SARS-CoV-2 RBD were detected with an indirect RBD-specific ELISA. Secondary antibody was the anti-human IgG (1:10,000) (Rockland Immunochemicals, USA). ROC analysis determined the positive OD_450_ cutoff value (dashed line). Positive plasma samples were donated COVID recovered patients and pre-pandemic serum samples were the negative controls. Based on the experimentally determined cutoff value, 64 of the 66 positive samples were anti-RBD positive, giving a sensitivity value of 96.96%. All but one of the negative samples were below the cutoff value for a specificity of 95.45%. Adjacent tables list first and third quartiles along with mean and median OD_450_ values of COVID-positive and negative human samples. Bars represent mean and standard deviation (n>3). (C) Anti-6His western blot on HEK-293T17 purified RBD from 5% serum conditions. Samples were run on a 12% denaturing SDS-PAGE gel. Protein was transferred to nitrocellulose and probed with anti-6His antibody at 1:10,000 (Proteintech, USA). White light and chemiluminescent images were overlaid and from left to right, ladder (lane 1) and purified RBD (lane 2). (D) Silver stain of purified recombinant SARS-Cov-2 RBD produced in HEK-293T17. From left to right: Ladder, HEK-RBD under 5% serum conditions, HEK-RBD from 2% serum conditions. Samples were denatured and run on a 12% SDS-PAGE gel and Silver Stained (Thermo Fisher Scientific, USA). For B and C, representative data shown.

To establish a baseline for future SARS-CoV-2 screening of companion animals, 128 pre-pandemic feline serum samples collected prior to December 2019 were retrospectively screened using our in-house ELISA. Nineteen samples were of a known FCoV serological status, with the remaining 109 of unknown FCoV status. Following the same protocol used for screening human serum samples (Fig. 1), feline samples were tested for antibodies against SARS-CoV-2’s RBD (Fig 2A). There were two batches tested. Serum samples from feral cats in East Tennessee collected from 2007 to 2012 (2007-2012) (n=36) and convenience samples from client-owned cats undergoing routine blood work (listed as Pre-pandemic) (n=92) (Fig 2A). As expected, SARS-CoV-2 experimentally infected cats [14] tested positive with high relative OD_450_, and normal cat serum (i.e., negative control) with very low relative OD_450_ (Fig 2A). Despite pre-dating the pandemic, 52% (67/128) of the cat samples tested positive for antibodies against SARS-CoV-2 RBD. This is surprising as there was a lack of high cross-reactivity in guinea pigs immunized with FCoV in Fig 1A. Several reports also showed a lack of similarity and cross reactivity between alpha coronaviruses and SARS-CoVs [18, 32, 38, 56]. Indeed, two other groups found that pre-existing immunity to FCoV had no impact on seropositivity of feline samples [38, 58]. To ensure that the positive ELISA results were specific to the RBD and not to a co-purified protein, a western blot was carried out using serum from a positive sample (Fig 2B). Positive cat serum bound a ∼32 kDa protein, the size of the RBD protein (Fig 1C). Notably, normal cat serum did not react with any other protein despite the presence of co-purified proteins. To further show the specificity of the anti-RBD response, we titrated seropositive and seronegative samples. Starting with serum from cats experimentally infected with SARS-CoV-2 (Fig 2A) and normal cat serum, saw a normalized OD_450_ >3 standard deviations above the negative control (i.e., normal cat serum) at all dilutions. This gives a titer >8100 (Fig 2C). 17 seropositive and 10 seronegative pre-2020 cat samples were titrated and assayed in our ELISA (Fig 2D). Titers ranged from 900 to 8100, with a median titer of 2700 demonstrating both a high anti-SARS RBD prevalence and titer. Titrations of these earlier seropositive and seronegative feral cat samples have a similarly high titer (Fig 2E) (median titer 8100) which is on par with the pre-pandemic samples. AUC for all groups is shown to the right side of their respective titration. AUC analysis of the titrated samples showed a significant difference between all positive and negative samples.

**Figure 2:**
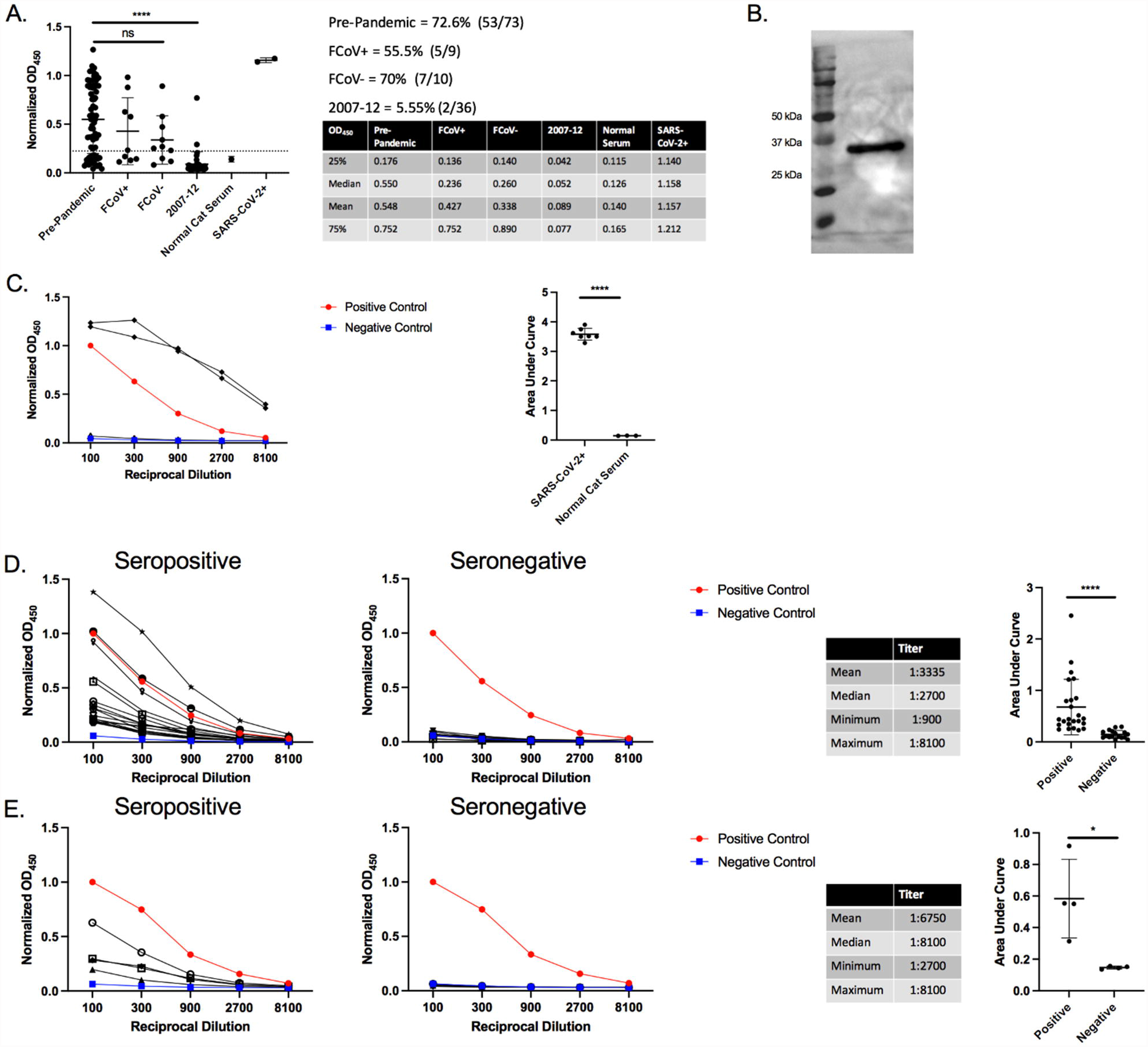
Pre-Pandemic Feline Antibodies Cross-React with SARS-CoV-2 RBD. (A) ELISA results of cat serum RBD reactivity. 93 pre-pandemic feline serum samples were tested for reactivity in our anti-RBD ELISA with anti-felid IgG secondary (1:10,000) (Invitrogen, USA). Cutoff values were determined by receiver operator curve (ROC) analysis. OD_450_ for samples in each group were plotted with the dotted line representing the positive threshold. Two sets of pre-pandemic cat samples were collected. Pre-pandemic cat convenience samples (n=73) were collected in local clinics and sent to the University of Tennessee for diagnostic testing or during feral cat studies (2007-2012) (n=36). Pre-pandemic convenience samples were subdivided into feline coronavirus positive (FCoV+) and negative (FCoV-) subgroups. Normal cat serum (Jackson ImmunoResearch Laboratories, USA) serves as the negative control and SARS-CoV-2+ serum from two cats experimentally inoculated with SARS-CoV-2 as positive controls. Side table lists first and third quartiles and mean and median OD_450_ values for all samples. Bars represent mean +/- standard deviation (n>3 for all samples). (B) Western blot of purified RBD using serum from a single positive cat sample. Purified RBD was run under denaturing conditions and blotted onto nitrocellulose. The RBD blot was first probed with cat serum from an ELISA positive sample (1:20 dilution) followed by anti-felid IgG-HRP conjugated (1:10,000 dilution) (Invitrogen, USA). White light and chemiluminescent images were overlaid. Lane 1 is the molecular weight ladder and lane 2 is purified RBD. (C, D, E) Titration of seropositive and seronegative serums assessed *via* RBD ELISA. OD_450_ values were plotted against the reciprocal dilution. Samples were considered positive if they were 3 standard deviations above the negative average for each dilution. Anti-RBD titer was designated as the last dilution above the negative cutoff. Positive controls were human COVID-positive serum and negative controls were normal human and cat serum (Jackson ImmunoResearch Laboratories, USA). Statistics for the positive sample titrations are included in the table along with AUC analysis. (C) Serum from two SARS-CoV-2 infected cats 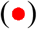 and normal cat serum 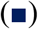 were titrated in an anti-RBD ELISA. (D) Titration of 17 seropositive and 10 seronegative, pre-pandemic cat samples. (E) Titration of four seropositive and seronegative cat samples collected from 2007-12. For A and B, representative data shown. For A, Tukey’s one-way ANOVA with multiple comparisons was performed. For C, D, E AUC analysis and Student’s one-tailed t-test with Welch’s correction was performed. p< 0.05 = *, p< 0.01 = **, p< 0.001 = ***.

Following the surprising presence and prevalence of anti-RBD responses in pre-pandemic cats, we explored the epidemiological characteristics of our samples. Pre-pandemic convenience samples were submitted to the University of Tennessee for diagnostic testing of feline herpesvirus, feline calicivirus, and FCoV. Age, sex, and location of seropositive and seronegative samples are shown in Table 1. Both seropositive and seronegative samples had a mean age of >3 years with no difference between the groups and contained similar ratios of male: female animals (Table 1). Seropositive samples were found in disparate geographic locations from opposite coasts of the United States (i.e., New York to California (Table 1)). This observation indicates that seropositivity is not confined to a single geographic region (e.g., East Tennessee). Based on our limited sampling, we were unable to identify any unique characteristic or identifier for seropositive vs seronegative samples.

With our discovery of pre-existing antibodies against SARS-CoV-2’s RBD, it was pertinent to examine samples from dogs, another companion animal with high human contact. Serum samples from dogs (n=36) were collected and retrospectively screened as part of a tick study during a 7-month period beginning in Jan 2020 and extending into July 2020. These samples are considered post-pandemic because the timeframe straddles the arrival of SARS-CoV-2 in East Tennessee (∼March 2020). The initial ELISA screen identified 97% seropositivity in the dog samples (Fig 3A) with only 1 sample falling below the cutoff established on human serum. Surprisingly, serum from purpose-bred research animals housed at the University of Tennessee also showed high levels of reactivity (Fig 3A). This raised suspicion about the specificity of the response. To address this, western blot analysis with canine serum (Fig 3B) identified a protein other than the RBD (see the ∼32 kDa protein in Fig 1C and 2B). The canine serum recognized a ∼60 kDa protein which is likely a co-purified protein present after RBD purification and is faintly seen in the silver-stained gel in Fig 1D. This co-purified protein was not detected in the blots performed for Fig. 1C and 2B using anti-6His monoclonal antibody and cat serum, respectively. Although there is a possibility that canine serum recognizes an oligomer of RBD [63], based on Fig 1C the anti-6His antibody does not detect any protein >32kDa, eliminating this possibility.

**Figure 3:**
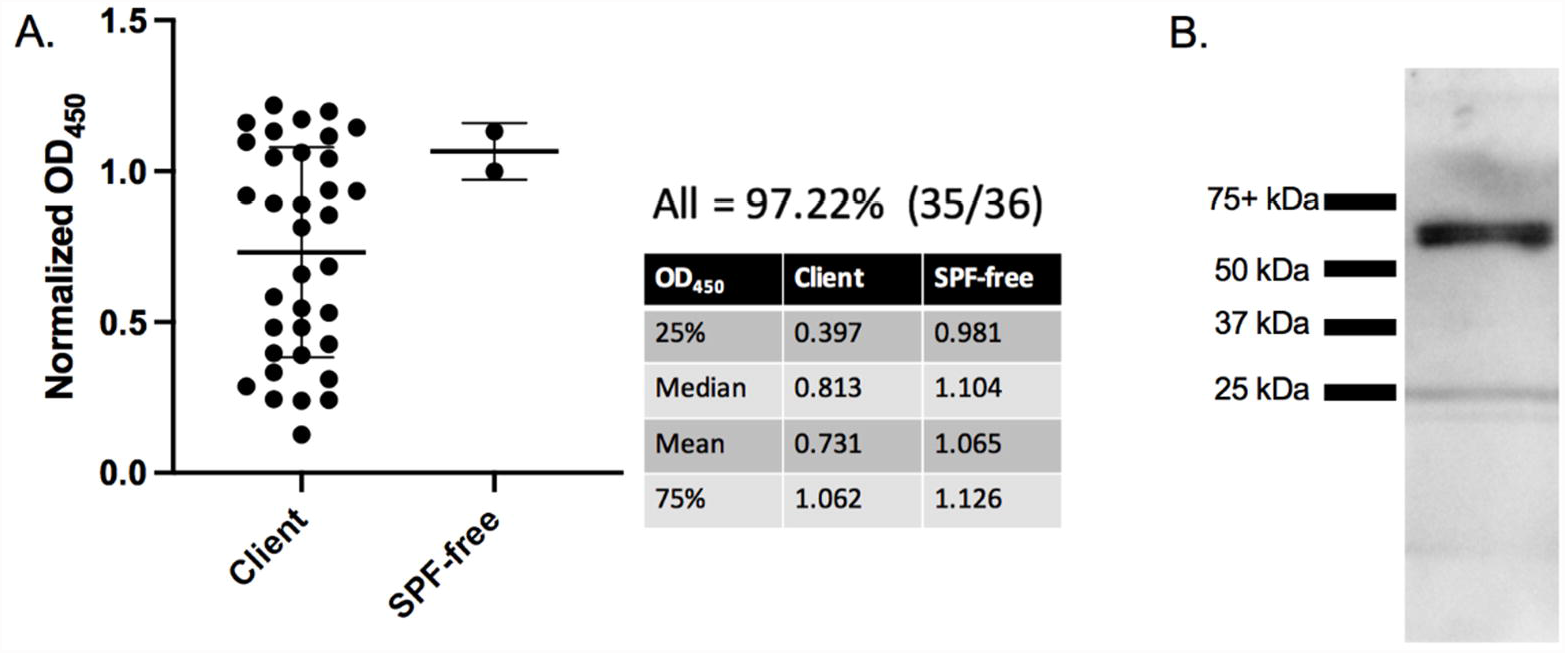
Dog Serum Cross-reacts to a Co-Purified Protein. (A) Anti-SARS-CoV-2 RBD ELISA with dog serum. Serum from thirty-six client-owned and two purpose-bred research dogs were tested in an anti-RBD ELISA with anti-canine IgG secondary HRP (1:10,000) (Bethyl Laboratories, USA). Table to the right lists the first and third quartiles, median, and mean OD_450_ values for all samples. Bars represent mean +/- standard deviation (n>3). (B) Western blot of purified RBD using serum from a positive dog sample. Purified RBD was probed with dog serum from an ELISA positive sample (1:20 dilution) followed by anti-canine IgG-HRP (1:10,000 dilution) (Bethyl Laboratories, USA). White light and chemiluminescent images were overlaid. Lane 1 (from left to right) ladder and lane 2: purified RBD. For all figures, representative data shown.

Following our observation of high levels of anti-SARS-CoV-2 RBD antibodies in North American cats, we began examining other regional animals. Serum from Tennessee resident, pre-pandemic cows (n=33) and tigers (n=9), post-pandemic East Tennessee elk (n=12), and post-pandemic South Carolina white-tailed deer (n=22) were tested for anti-SARS-CoV-2 RBD antibodies (Fig 4A). Of the four species tested, only the deer from South Carolina showed any seropositive samples (2/22). Serum titrations show the two seropositive samples have a high titer >8100 (Fig 4B), and AUC of the titrations show a significant difference between seropositive and negative deer samples (Fig 4C). Unfortunately, due to limited sample volume, we were unable to run western blots to demonstrate the specificity for the RBD protein. The deer are post-pandemic and could represent recent transmission of SARS-CoV-2 into the deer population. Although these animals probably have had limited contact with humans, white-tailed deer are susceptible to and capable of transmitting SARS-CoV-2 [64]. Another possibility is that this species was exposed to the same etiological agent as our pre-pandemic seropositive cats.

**Figure 4:**
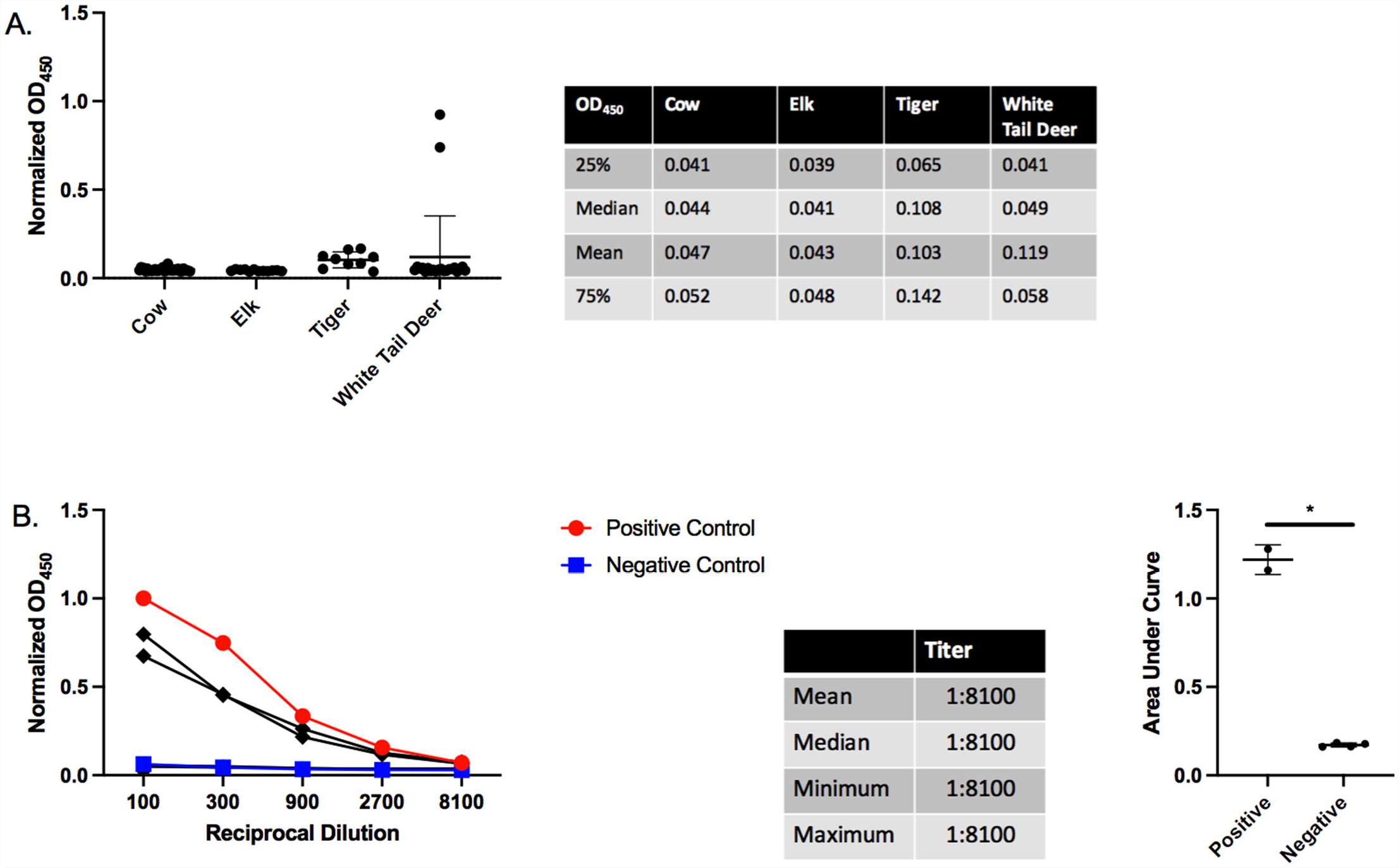
Serological Testing of Other Regional Animals. (A). Anti-SARS-CoV-2 RBD ELISA with bovine, elk, tiger, and deer serum. Thirty-three pre-pandemic East Tennessee cows, twelve post-pandemic East Tennessee elk, nine pre-pandemic East Tennessee tigers, and twenty-two post-pandemic South Carolina deer serum samples were tested for anti-RBD antibodies. Species-specific secondary antibodies were used at the following dilutions: anti-bovine 1:250 (Bethyl Laboratories, USA), anti-elk/deer 1:250 (KPL, USA), anti-tiger/cat 1:10,000 (Invitrogen, USA), and anti-deer 1:250 (KPL, USA). Bars represent mean +/- standard deviation (n>3 for all samples). (B) Titration of two seropositive 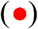 and four seronegative 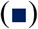 deer samples. OD_450_ values are plotted against the reciprocal dilution of each sample. Samples were considered positive if they were 3 standard deviations above the negative average for each dilution. Positive and negative controls were human COVID-positive and negative samples, respectively. Statistics for the positive sample titrations are included in the table. The AUC analysis for titrations of deer ELISA positive and negative samples is shown to the right. For all figures, representative data shown. For AUC analysis Student’s one-tailed t-test with Welch’s correction was performed. p< 0.05 = *, p< 0.01 = **, p< 0.001 = ***.

To address whether our ELISA-positive animal samples can neutralize SARS-CoV-2 infections, we measured the ability of cat serum to block the interaction of the spike protein with the human ACE-2 (hACE-2) receptor using a commercially available flow cytometry-based bead assay. In this assay, neutralization is characterized as the decrease in fluorescence when antibodies block the fluorescently labeled SARS-CoV-2 S1 subunit from binding to hACE-2 conjugated beads (Fig 5A). Because this assay is not species specific or immunoglobulin type dependent, it is applicable for assessing both human and feline serum. The internal antibody control shows a decrease in fluorescence corresponding to levels of neutralizing monoclonal antibody against SARS-CoV-2. Serum from experimentally infected cats showed potent neutralization at 1:100 dilution. However, only one ELISA-positive, pre-pandemic cat sample showed neutralization (Fig 5B). One of the seropositive white-tailed deer samples, and a single serum sample from mice immunized with PRCoV also showed slight neutralization, clearing the determined ROC threshold/cutoff value (Fig 5B). Notably, we were unable to detect high levels of neutralization/neutralizing antibodies even in several of the human convalescent serum samples (Fig 5B).

**Figure 5:**
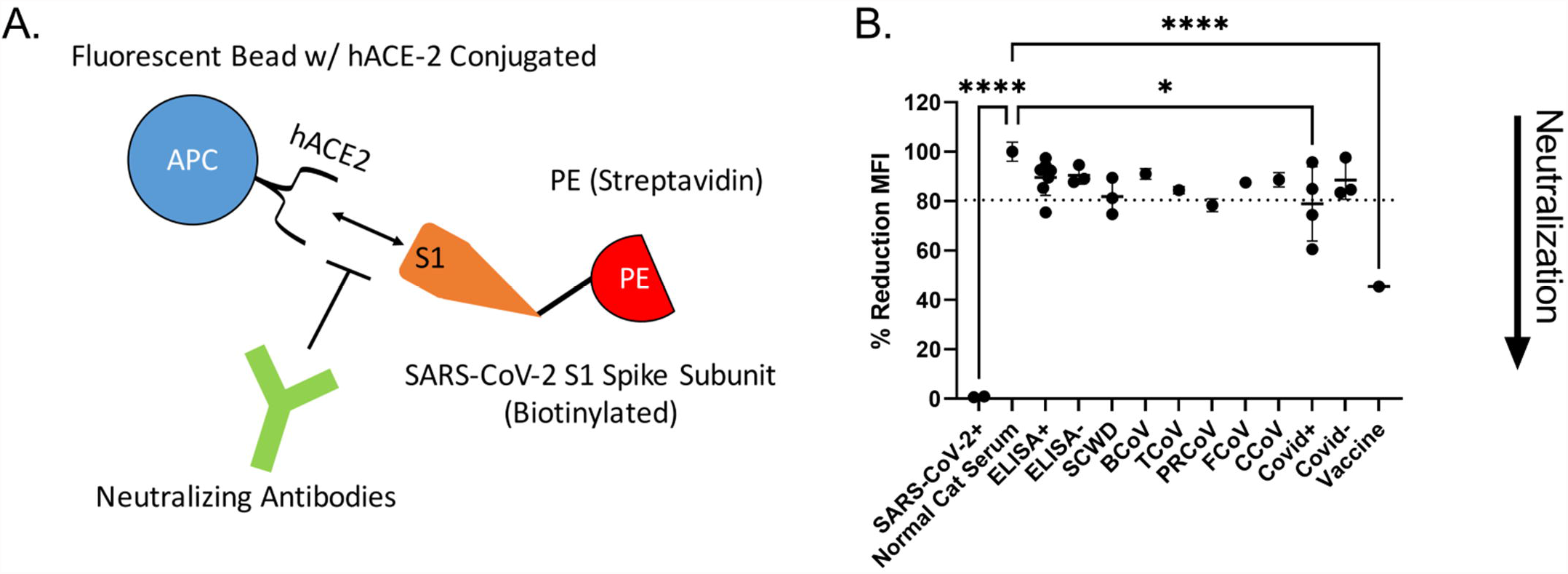
Neutralization Assays. (A) Schematic of the neutralization assay. Neutralization is measured as the decrease in binding of phycoerythrin (PE)-labeled SARS-CoV-2 S1 subunit to human ACE-2 conjugated beads. Addition of neutralizing antibodies results in a decreased mean fluorescent intensity (MFI) as measured by flow cytometry. (B) Neutralization of SARS-CoV-2 S1 subunit interaction with hACE2. Serum from several ELISA positive and negative cats (ELISA+ and ELISA-, respectively), serum from South Carolina white-tailed deer (SCWD, 2 ELISA-positive and 1 negative), mice immunized with other common coronaviruses (BCoV=bovine coronavirus, TCoV= Turkey coronavirus, PRCoV=porcine respiratory coronavirus, FCoV=feline coronavirus, CCoV=canine coronavirus), and human serum samples (Covid+ = convalescent plasma from Covid+ humans, Covid- = pre-SARS-CoV-2 serum samples, Vaccine = serum post vaccination against SARS-CoV-2 Spike protein). SARS-CoV-2 infected cats and normal cat serum served as positive and negative controls, respectively. Data was normalized to normal cat serum representing 100% binding of SARS-CoV-2 S1 subunit to hACE2 beads. ROC analysis was used to generate a positive reduction threshold (dotted line). Each point is an average of 2 replicates. A Tukey’s one-way ANOVA with multiple comparisons was used to analyze experimental groups. p< 0.05 = *, p< 0.0001 = ****.

Because cross-reactivity of antibodies to SARS-CoV-2 RBD independent of SARS-CoV-2 infection has not been previously reported in felines, we suspected that the etiological agent could be another coronavirus [38, 58]. Fecal samples were collected from healthy East Tennessee cats and screened for coronaviruses using pan-coronavirus primers amplifying conserved regions of the RNA-dependent RNA polymerase (RdRp), helicase (Hel), and spike (S) genes [60]. Coronavirus viral RNA, whether common animal coronavirus or SARS-like coronavirus, is potentially shed in feces [8, 9, 11, 13]. Collection of fecal samples represented a non-invasive collection method, and SARS-CoV-2 has been reported to have prolonged shedding within fecal samples of humans [8, 9, 11, 13]. Fifteen out of thirty samples (50%) tested positive for at least one loci, with most yielding positive results for multiple loci (Table 2). Not surprisingly, sequences cluster within the alpha-coronavirus group and with high similarity to previously identified FCoV strains. When all five loci were aligned and concatenated together, the Maximum-Likelihood phylogenetic tree places the concatenated coronavirus sequences within the alpha-coronavirus lineage, closely related to FCoV (Fig 6A). We were unable to amplify or identify any sequences which resemble SARS-like coronaviruses or beta-coronaviruses. Partial sequencing of the S1 region was able to amplify the RBD from several coronavirus RNA positive samples. The sequenced RBDs were again highly similar to FCoV based on Maximum-Likelihood phylogenetic tree (Fig 7A). Along with the phylogenetic tree, a similarity matrix demonstrates high RBD similarity between previous FCoV strains and those sequenced here (∼80%) (Fig 7B). The RBD from these fecal samples displays low similarity to betacoronaviruses such as MERS, SARS-CoV, and SARS-CoV-2 (∼30%) as previously reported (Fig 7B).

**Figure 6:**
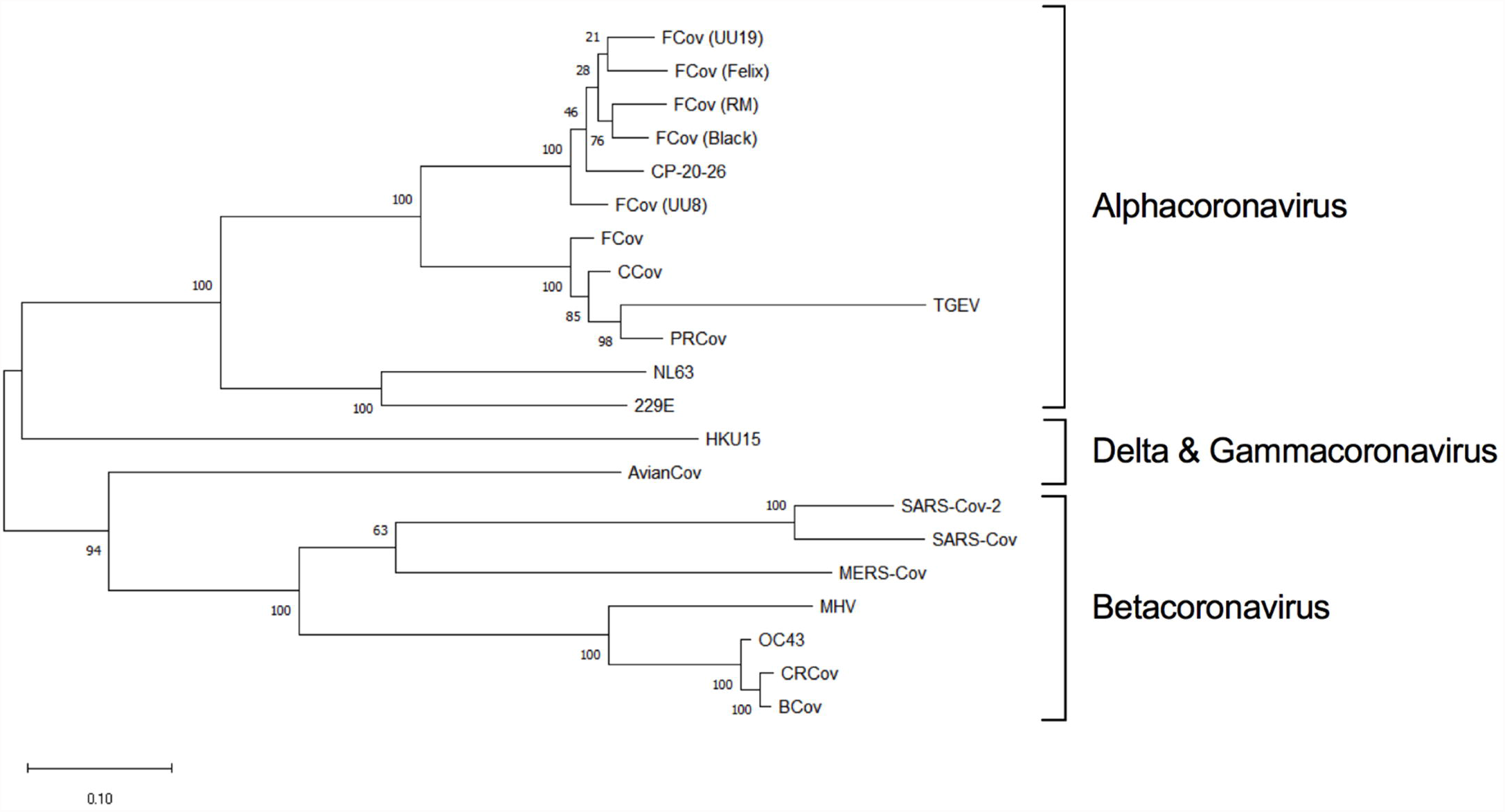
Pan Coronavirus Screen of East Tennessee Felines. Fecal samples from healthy cats were collected and screened for conserved coronavirus sequences. Phylogenetic tree were generated consisting of common human and animal coronaviruses: CRCoV (canine respiratory coronavirus), BCoV (bovine coronavirus), OC43 (human beta-coronavirus), MHV (murine hepatitis virus), MERS-CoV (Middle East respiratory coronavirus), SARS-CoV-2 (severe acute respiratory coronavirus-2), SARS-CoV (severe acute respiratory coronavirus), AvianCoV (duck coronavirus), NL63 (human alpha-coronavirus), 229E (human alpha-coronavirus), TGEV (transmissible gastroenteritis virus), PRCoV (porcine respiratory coronavirus), FCoV (feline coronavirus strains UU19, Felix, RM, Black, UU8), CCov (canine coronavirus), HKU15 (porcine delta-coronavirus), as well as a locally identified coronavirus (CP-20-26). Sequences from five coronavirus loci were independently aligned, trimmed, and concatenated together. Concatenated sequences were aligned, and phylogenetic trees generated with the Maximum-Likelihood method with bootstrap analysis in MEGA X. Bootstrap values for each branch are shown with lengths to scale. Coronavirus lineages are annotated on the tree.

**Figure 7:**
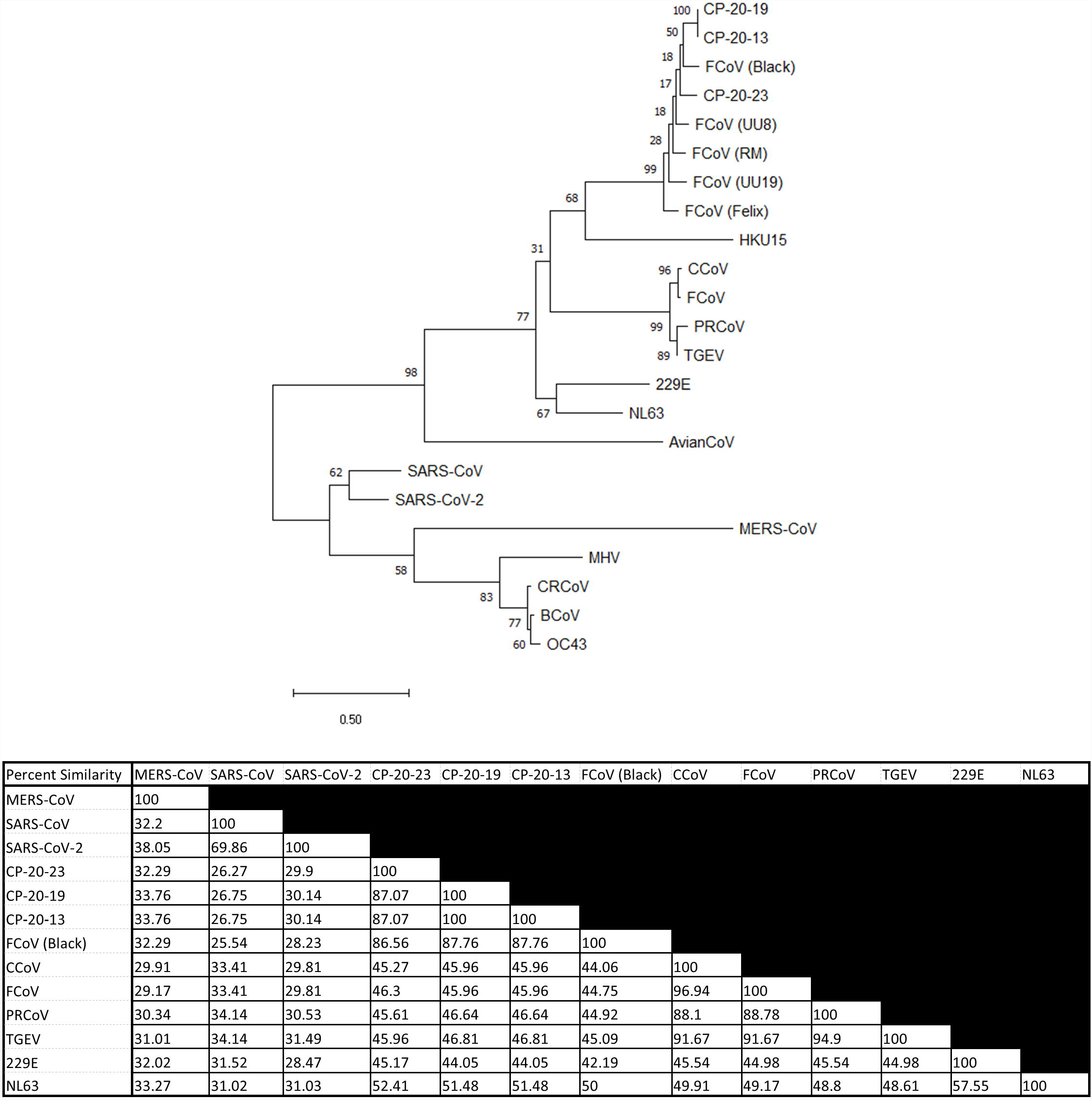
RBD Coronavirus Screen of East Tennessee Felines. (A) Fecal samples from healthy cats were collected and screened for S1/RBD coronavirus sequences. Phylogenetic tree consisting of common human and animal coronaviruses: CRCoV (canine respiratory coronavirus), BCoV (bovine coronavirus), OC43 (human beta-coronavirus), MHV (murine hepatitis virus), MERS-CoV (Middle East respiratory coronavirus), SARS-CoV-2 (severe acute respiratory coronavirus-2), SARS-CoV (severe acute respiratory coronavirus), AvianCoV (duck coronavirus), NL63 (human alpha-coronavirus), 229E (human alpha-coronavirus), TGEV (transmissible gastroenteritis virus), PRCoV (porcine respiratory coronavirus), FCoV (feline coronavirus strains UU19, Felix, RM, Black, UU8), CCoV (canine coronavirus), HKU15 (porcine delta-coronavirus), as well as locally identified coronaviruses (CP-20-13, CP-20-19, CP-20-23). Sequences from the S1 region were aligned and trimmed. Maximum-Likelihood phylogenetic trees were generated with bootstrap analysis in MEGA X. Bootstrap values for each branch are shown with lengths to scale. (B) Percent similarity matrix of select coronaviruses. Aligned and truncated RBD regions from the shown coronaviruses were analyzed *via* Clustal Omega to determine percent identity matrix.

## Discussion

We have employed serological screening as a method to detect potential SARS-CoV-2 exposure in animal populations. Tracking active viral spread in wild and domesticated animals in real-time *via* sequencing and RT-qPCR is expensive and low throughput. In addition, the unknown transient nature of viral shedding from different secretions/locations makes this type of surveillance prohibitively expensive with no guarantee of identifying a virus. Serological detection of antibodies against SARS-CoV-2 is comparatively high-throughput and inexpensive while still maintaining sensitivity. A downside of this methodology is the lack of an up-to-date picture of cross-species transmission, as serology trails initial infections by several days to weeks [22, 33]. On the other hand, due to the lowered cost of serological testing there is a compensatory increase in testing capability allowing a broader swath of animals and regions to be sampled with more frequent re-sampling to track spillover into new species. Our adapted protocol yields recombinant SARS-CoV-2 RBD protein allowing production of a low-cost indirect anti-RBD ELISA. The recombinant RBD was sufficient for serological screening *via* ELISA and is amenable to most labs with prior tissue culture capabilities and does not require large initial investments in cell lines, culture media, or specialized incubators. We validated our method demonstrating low cross-reactivity with other common animal coronaviruses (Fig 1A) and >95% sensitivity and specificity on human serum samples (Fig 1B).

For the seropositive samples identified in our study, mean titers for positive cat samples were relatively high at ∼2700 (Fig 2E, G), which based on reported rapid declines in anti-RBD responses for SARS-CoV and FCoV points to exposure within the past few years [37, 38, 40]. Further, based on FCoV studies, animals with high titers typically correlate with active viral shedding and spread within a household, which highlights a potential overlap between seropositivity and viral shedding [37]. This is in stark contrast to SARS-CoV-2 serosurveys on pre- and post-pandemic feline samples from Central China. They found no evidence of exposure before the outbreak, but also positivity levels post-pandemic were significantly lower than shown here (∼12% Central China vs >50% USA) (Fig 2A) [38, 58]. OD_450_ and titers of pre-pandemic seropositive cat samples, while high, were lower than SARS-CoV-2 experimentally inoculated cats (6 weeks post infection) (Fig 2A, C, E, G). This likely represents a natural decline in titer over time for the environmental samples but could also represent lower titers of cross-reactive antibodies from another coronavirus.

Unfortunately, dog serum was shown to bind to a co-purified protein (Fig 3B), leaving us unable to utilize our assay for examining cross-species transmission of SARS-like coronaviruses to canines. We can show that recombinant RBDs produced and purified by groups at both Mt. Sinai and Emory contain co-purified proteins at approximately the same size as shown in Fig 1A [30, 33]. As such, screens for SARS-CoV-2 exposure in canines would likely require producing and purifying the RBD using a different strategy that eliminates non-RBD protein contamination. Recently both SARS-CoV-2’s RBD and soluble full-length spike have been produced and purified in plants [65]. This alternative method may prove useful for animal SARS-CoV-2 screening by reducing or eliminating false positives due to co-purified proteins.

After the discovery of pre-pandemic seropositive cats, we examined other commercial (cow) and convenience samples from local wild species (deer, elk, tiger) (Fig 4A). Two out of twenty-two (9%) white-tailed deer from South Carolina were positive for antibodies against the RBD (Fig 4B). Unlike our cat samples, the two seropositive deer could represent transmission of SARS-CoV-2 into the local deer population because these samples were collected post-pandemic. Interestingly, a recent report showed that ∼40% of white-tailed deer from 4 states (Illinois, Michigan, New York, and Pennsylvania) were positive for SARS-CoV-2 antibodies [29]. Seropositive animals were only observed from 2019 onward, with pre-pandemic deer testing negative on their SARS-CoV-2 neutralization assay. This information supports our finding in South Carolina deer (Fig 4). SARS-CoV-2 sequences were also recently isolated from the retropharyngeal lymph nodes of wild and captive deer [27, 28]. The dominant genotype of deer-isolated SARS-CoV-2 genotypes closely corresponded to those circulating within humans at the time, pointing to potential rapid transmission from humans to animals [28]. This highlights the importance of the One Health Initiative to provide information on the potential exposure and spillover into other species and whether there is recombination with native coronaviruses occurring to generate new variants or establishment of new reservoirs in North America. Further work is needed to determine the prevalence, spread, and identity of SARS and other coronaviruses circulating within North American deer and associated species.

That samples from cats experimentally infected with SARS-CoV-2 displayed potent neutralization (Fig 5B) is unsurprising because of their high ELISA titers (Fig 2C). These samples were collected at ∼8 weeks post infection and likely represent peak titer and neutralization capacity [14]. Neutralization of SARS-CoV-2 S1 subunit was variable for environmental feline ELISA-positive samples (Fig 5B). There was no significant difference in MFI/neutralization between the anti-RBD seropositive and seronegative feline samples (Fig 5B). Even sera from convalescent, COVID-recovered individuals showed none to minimal neutralization (Fig 5B). Several groups have found that not all anti-RBD responses generate neutralizing antibodies [50-53, 66]. Indeed, even in convalescent serum, high levels of RBD recognition does not guarantee high neutralizing titers, consistent with our own observations (Fig 5B) [50]. Based on ELISA and neutralization results (Fig 2A, Fig 5B), we suspect that these animals contain antibodies recognizing SARS-CoV-2’s RBD, but likely bind to non-neutralizing epitopes of the RBD domain.

Because ∼50% of cats surveyed were seropositive, we reasoned that isolation of the suspected infectious agent or coronavirus might be possible. Based on fecal viral RNA shedding following animal coronaviruses infections, PCR amplification with universal coronavirus primers was used to screen for potential causative agents of anti-RBD seropositivity. This allowed for non-invasive testing and isolation of coronavirus RNA from infected cats. In line with previous studies on other wild animals, we did not identify any non-alphacoronaviruses circulating in felines [67-70]. The coronavirus sequences that were isolated and sequenced likely represent normal circulating FCoVs (Fig 6). Due to the opportunistic nature of our sampling, we were unable to obtain any paired blood and fecal samples from the same animal. As such, we are unable to conclude whether the cats with identified FCoVs would produce cross-reactive antibodies against SARS-CoV-2’s RBD. However, based on our ELISA results in Fig 1A and 2A showing no correlation with SARS RBD antibodies and FCoV infection, we would suspect not. Furthermore, not knowing when the seropositive cats were exposed (i.e., cats could have been infected potentially years prior to any fecal sampling) fecal sampling and sequencing would not detect a novel coronavirus if it had been cleared. Following the identification of coronavirus positive fecal samples, we attempted to amplify and sequence the entire RBD region from positive samples to look for similarity to SARS-CoV-2 or SARS-like viruses. Large portions of the S1 region spanning the RBD were sequenced and contained RBD regions similar to previously isolated FCoV strains (Fig. 7), with no similarity to SARS or betacoronaviruses.

The current study demonstrates cross-reactivity of pre-pandemic feline samples with the RBD of SARS-CoV-2. Our indirect ELISA screen has provided evidence for seropositivity of serum from North American cats and deer to a SARS-CoV-2 protein previously shown to be highly specific to SARS coronaviruses [32, 38, 46, 49]. This is the first study to demonstrate seropositivity of animal samples pre-pandemic. What induces this cross-reactive response was not readily apparent. However, we propose several possibilities: exposure to another infectious agent generating cross-reactive antibodies, infection with multiple common coronavirus strains (Feline coronavirus or otherwise) generating cross-reactive antibodies, or exposure of animals to a SARS-like coronavirus pre-pandemic.

There is evidence both for and against these explanations for the seropositivity observed. While we cannot discount a non-coronavirus infection generating cross-reactive antibodies, the RBD of SARS-like viruses is thought to be unique with no previous evidence of RBD cross-reactivity [32, 38, 46, 49]. A plausible explanation for seropositivity against SARS-CoV-2’s RBD is infections with coronaviruses generating an atypical response. To-date, cross-reactivity against the RBD of SARS-CoV-2 has only been reported for SARS and SARS-like coronaviruses [32, 46]. The common human and animal coronaviruses (both alpha and beta coronavirus families) individually do not generate cross-reactive antibodies against this protein, presumably making it a SARS-specific response [30, 32, 33, 38, 46, 49]. For example, prior FCoV exposure did not impact SARS-CoV-2 RBD recognition [38]. Our own observations further demonstrated that FCoV (both serotype I and II) and TGEV did not correlate with serostatus for SARS-CoV-2’s RBD (Fig 2A) (FCoV+, FCoV-). Additionally, humans and animals are exposed to multiple coronaviruses throughout their lifetime, generating measurable immune responses to antigenic proteins [71, 72]. With humans, CoV infections occur on average once a year, with protective immunity appearing to wane a few months after initial exposure [71, 72]. Despite reports of cross-reactivity towards portions of SARS-CoV-2’s spike protein induced by other human coronaviruses, there have been no reports of cross-reaction against the distinct RBD region [32, 46, 49]. While it does not disprove multiple CoV infections generating cross reactive antibodies, it does cast doubt on this possibility. This leads us to propose prior exposure to a SARS-like coronavirus for North American cats.

There is limited evidence to support prior transmission of a SARS-like virus within felines. However, cats are susceptible to infection and transmission of SARS-CoV-2 [14]. SARS coronaviruses evolved from bat coronaviruses and maintain a high degree of sequence similarity, even in the RBD region [32]. Conceivably, our definition of SARS-like coronavirus could also encompass bat coronaviruses due to amino acid similarity and potential cross-reactivity in the RBD region. At least within Tennessee there are numerous cave systems with several native species of bats potentially leading to an inter-species transmission. Feline and bat interactions could lead to direct transfer of a SARS-like agent, or an intermediary species could be involved. North American deer mice were recently shown to be susceptible to human SARS-CoV-2 and capable of mouse-to-mouse transmission representing another potential point of introduction into the feline population [73]. The pre-pandemic exposure of cats to a SARS-like agent does have detractions. Other groups examining pre-pandemic samples have failed to find evidence of RBD-reactive serum even within Central China [38, 56, 58]. The ELISA-positive samples from our environmental samples (i.e., feline and deer) did not neutralize to the same degree as experimentally inoculated felines (Fig 5B) [14]. However, significant neutralization was not observed for most samples, even convalescent serum from COVID-positive individuals (Fig 5B). Despite ∼50% seroprevalence in our samples, we were unable to identify any coronavirus sequences capable of eliciting an anti-SARS RBD response (Fig 6 and 7). Additionally, there is currently no evidence of a circulating SARS-like or bat betacoronavirus in North America [67-70]. While we cannot conclusively demonstrate the origin of the anti-RBD responses shown here, this study indicates that there could be a virus (or another infectious agent) that can generate cross reactive antibodies to the SARS-CoV-2 RBD.

## Limitations of this study

A major limitation of this study is the number of serum samples surveyed for all species. Due to the small sample size, we have refrained from making extrapolations from our data to larger animal populations or geographic regions. Similarly, a relatively small number of feline fecal samples were screened for coronavirus viral RNA (n=30), limiting our ability to detect coronaviruses. Our choice of sampling source may have also limited our ability to detect novel coronaviruses or SARS-like coronaviruses. We chose to utilize fecal samples for coronavirus screening because it is non-invasive, animal coronavirus shedding within feces is common, and many are transmitted fecal to oral. Additionally, SARS-CoV and SARS-CoV-2 viral RNA has been detected in stool samples of infected humans [8, 9, 11, 13]. This fecally shed RNA persists longer than nasal or oral sources [13]. However, sampling nasal/oral sources rather than fecal samples may allow for better detection of SARS-CoV-2. Indeed, isolation of SARS-CoV-2 RNA and whole genome sequencing was possible from deer retropharyngeal lymph nodes [28]. Along with sampling site limitations, infection and shedding of viral RNA is a transient process with a relatively small window to detect and sequence the causative agent. While ∼20 feline samples were tested for anti-FCoV antibodies, the majority of our samples were of an unknown FCoV status. Additionally, medical histories were not available for these animals and were not screened for prior exposure to other infectious agents. As such, we cannot definitively rule out some other infectious agent conferring the anti-RBD responses.

## Conclusion

Our initial goal was to develop an ELISA assay for tracking the reverse zoonosis of SARS-CoV-2. However, when establishing our baseline on pre-pandemic cat samples we discovered seropositive serum for an antigen previously reported to be SARS-specific (i.e., RBD). Seropositivity was ∼50% in feline samples and could be found several years prior to the genesis of the current SARS-CoV-2 pandemic. What generated the RBD-reactive antibodies is unknown, but we proposed three possibilities: cross-reaction caused by another infectious agent, multiple coronavirus infections creating a rare cross-reactive antibody, or existence and circulation of a SARS-related virus containing the RBD sequence. Regardless, the high rate of SARS-CoV-2 RBD seropositivity within a common companion animal further highlights our need for a better understanding of the prevalence and crossover potential of wild coronaviruses. Further investigations should address shedding of viral RNA from the seropositive species (i.e., cats and deer) identified here to isolate, sequence, and identify the agent enabling cross-reactivity against the RBD of SARS-CoV-2.

## Acknowledgments

We would like to thank the following for their generous contributions making this work possible: MEDIC Regional Blood Center

Mark Slifka – Oregon Health Sciences University

Jon Wall and Steve Foster – University of Tennessee Medical Center Knoxville

Krammer Lab – Icahn School of Medicine at Mount Sinai

Wrammert Lab – Emory University

Angela Bosco-Lauth – Colorado State University

Jennifer Weisent, Rebekah DeBolt, Sarah Englebert, Jessica Thompson – University of Tennessee College of Veterinary Medicine Shelter Medicine Service

